# Histone modifications and active gene expression are associated with enhanced CRISPR activity in de-silenced chromatin

**DOI:** 10.1101/228601

**Authors:** René Daer, Cassandra M. Barrett, Karmella A Haynes

## Abstract

Recently we demonstrated that closed chromatin composed of Polycomb proteins and histone 3 lysine 27 trimethylation impedes CRISPR-mediated genome editing by blocking the accessibility of chromosomal DNA to spCas9/sgRNA. Editing efficiencies were higher in cells where the same reporter locus had not been repressed, thus we presume that silenced chromatin can be modified to generate a Cas9-accessible state. To test this idea, we exposed the locus to antagonists of Polycomb silencing: Gal4-p65, a targeted transcriptional activator, and UNC1999, a chemical inhibitor of the histone H3K27 methyltransferase EZH2. For both we observed loss of histone trimethylation. Only Gal4-p65 treatment increased target gene expression. Initial Gal4-p65 overexpression impedes Cas9 activity, while a 9-day recovery period leads to enhanced Cas9 efficiency up to 1000 bp from the Gal4 binding site. No enhancement was observed with UNC1999. These results demonstrate the strong influence of transcription-driven chromatin remodeling on CRISPR editing at closed chromatin.

## INTRODUCTION

CRISPR/Cas9, a nucleic acid modifying system derived from prokaryotes, is a popular tool for rapid, high-throughput genome engineering in eukaryotic nuclei.^1,2^ Mounting evidence suggests that chromatin, the RNA-DNA-protein complex that mediates eukaryotic chromosomal organization and gene expression,^3–5^ inhibits Cas9 binding and nuclease activity *in vitro*^6–8^ and at many genomic sites in metazoan cells.^9,10^ While many Cas9 gene editing efforts have been successful, the complex structure of chromatin and its variation in different cells still pose a barrier to reliable and consistent use.^8–11^

A growing body of work has determined the strong influence of chromatin on the accessibility of DNA to Cas9, guide RNA complexes (Cas9/gRNA), and Cas9 editing efficiency.^8–11^ A series of *in vitro* studies using reconstituted nucleosomes formed from synthetic DNA and histones have demonstrated that Cas9-mediated binding and cleaving of nucleosomal DNA is completely blocked.^6–8^ Although these results might have limited relevance for chromosomal DNA since synthetic sequences used in these studies associate more tightly with histones,^12,13^ studies in mammalian cells corroborate the observation that chromatin structure can impede CRISPR activity. Surveys of CRISPR efficiencies at chromosomal sites in HEK293T, HeLa, human fibroblast^14^, and mouse embryonic stem cells^15^ have shown that heterochromatin (closed chromatin) is less accessible to CRISPR editing than euchromatin (open chromatin). Utilizing transgenic cell lines, we and others have shown that ectopic HP1-mediated^9^ and Polycomb Repressive Complex (PRC)-mediated^10^ heterochromatin decrease, and in some cases completely block, Cas9 binding and editing in HEK293 cells. Interestingly, editing at three out of nine sites within a PRC-repressed region was not blocked, suggesting that inhibition of Cas9 can be discontinuous across regions of closed chromatin.^10^ *In vivo*, dynamic chromatin organization poses a barrier to reliable CRISPR-mediated editing as demonstrated in zebrafish.16.

Research so far suggests that remodeling or disruption of closed chromatin might generate a more Cas9-permissive state. For instance, a recent study showed that deactivated Cas9 (dCas9) can be used as an an artificial pioneer factor to increase DNase hypersensitivity, which indicates depletion or relocation of bound proteins (i.e., histones), at several loci in mouse embryonic stem cells^17^. In another study, dCas9 was used to increase accessibility of a nearby target site to an active Cas9 nuclease^18^. This effect was was observed at sites within 50 bp of the dCas9 target. For CRISPR applications where unique pioneer factor target sites are not available at this proximity, more distal remodeling activity may be desired. To achieve greater control over Cas9-mediated editing in heterochromatin, further investigation is needed to determine the relationship between CRISPR activity and the impact of dynamic features found across broader regions, including transcriptional elongation, histone modifications, and association of non-histone proteins (Fig. 1).

**Figure 1.**
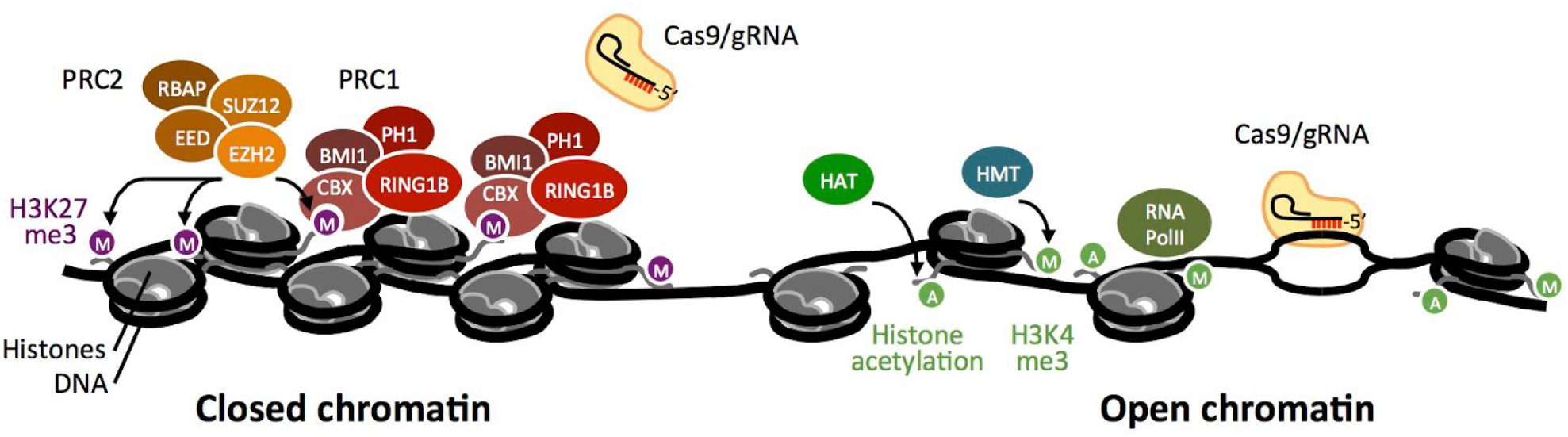
Recent studies suggest that facultative heterochromatin inhibits Cas9-mediated editing while open chromatin is permissive to Cas9. (A) The PRC2 (Polycomb repressive complex 2) holoenzyme generates the silencing mark histone 3 lysine 27 trimethylation (H3K27me3) (purple M). PRC2 includes Suppressor of Zeste 12 (SUZ12), Embryonic ectoderm development (EED), Retinoblastoma-binding protein (RbAp), and Enhancer of zeste 2 (EZH2).^3,5^ Polycomb repressive complex 1 (PRC1) includes Chromobox protein homolog (CBX), Ring finger protein 1b (RING1B), and Polycomb group RING finger protein 4 (BMI1).^3,5^ Polycomb proteins support histone compaction and block access of DNA to RNA polymerase and Cas9.^9,10^ Chromatin remodelers (not shown), histone acetyltransferases (HATs), and histone methyltransferases (HMTs), generate modifications that support open chromatin, accessible DNA, and a transcriptionally permissive state.^19,20^

To this end, we investigated how perturbations of the closed chromatin state impact Cas9-mediated editing by attempting to re-open chromatin that was previously closed by Polycomb-complex (PRC) formation at an ectopic site. Polycomb group (PcG) protein-driven gene silencing is conserved across metazoans, from simple echinoderms to plants and large land vertebrates^21^. PRC-mediated silencing (Fig. 1A) is elevated in stem cells ^22,23^ and cancer cells^24–27^ at loci that are central to cell development and are therefore likely targets of interest for gene editing. Polycomb-regulated targets are distributed over thousands of sites along chromosome arms in gene-rich regions. Therefore, PRC’s present a significant barrier to genome engineering efforts. In our previous work, we tested a variety of artificially-induced expression states at a PRC-repressed reporter gene and found that siRNA-mediated knockdown of the PcG protein Suz12 (Fig. 1) was accompanied by partial recovery of Cas9-mediated editing efficiency at a single CRISPR target site.^10^

Here, we expanded our investigation to determine transcriptional states and histone modifications that are associated with CRISPR-accessibility over a broader region of DNA (1000 bp). Experiments with transcription-activating proteins in cells^28^ and with nucleosome arrays *in vitro*^7,8^ suggest that expression-associated chromatin remodeling could improve Cas9 efficiency in human cells. To test this idea we measured two signatures of the conversion from epigenetic silencing to activation, gene expression and activation-associated histone modifications, at a PRC-repressed site that was subjected to transcription factor targeting or specific inhibition of H3K27 methylation. We determined the associated changes in CRISPR editing efficiency using a sensitive deep sequencing assay. We found that strong induction of transcription by the p65 subunit of NF-kB followed by culturing for several days resulted in sustained gene expression and the restoration of full Cas9 activity at a target site approximately 1000 bp downstream of the promoter. Specific inhibition of the histone K27 methyltransferase EZH2, and concomitant depletion of H3K27me3, was not sufficient to enhance gene expression nor Cas9 editing efficiency. Our results suggest that reorganization of chromatin after transcriptional elongation supports accessibility of DNA to Cas9 activity over a broad region of chromatin.

## MATERIALS AND METHODS

### Cell Lines and Plasmids

Development of the Luc14 and Gal4-EED/luc HEK293 cell lines is described in Hansen et al.^29^ Cas9/gRNA-expressing cells were generated via transfection of pU6-(BbsI)_CBh-Cas9-T2A-EGFP (DNASU UnSC00746685), which was built from pX330-U6-Chimeric_BB-CBh-hSpCas9 (a gift from Feng Zhang, Addgene plasmid #42230) as previously described^30^.

pU6-(BbsI)_CBh-Cas9-T2A-EGFP generates a single mRNA transcript that encodes the human optimized spCas9 protein and EGFP, separated by a T2A translation-skipping signal. Guide RNAs sg025, sg032, sg046, and sg048 were cloned into pU6-(BbsI)_CBh-Cas9-T2A-EGFP as previously described^30^. Gal4-p65 (a Gal4-mCherry-p65 fusion protein) was expressed from plasmid CMV-Gal4p65_MV1 (DNASU UnSC00746686). Annotated sequences for the plasmids used in this study are available online at https://benchling.com/hayneslab/f_/V1mVw1Lp-chromatin-crispr-interference/.

### Cell Culturing and Transfection

Cell culturing, *luciferase* silencing, and transfections of the Luc14 and GAL4-EED HEK293 cell lines were performed as previously described^30^. Gal4-EED/luc cells were treated with doxycycline (dox) for 96 h to induce silencing, then cells were grown in fresh dox-free medium (DMEM, 10% fetal bovine serum, 5% penicillin-streptomycin) for 24 h. For Gal4-p65 overexpression, ∼2.0E5 cells per well were transfected in 12-well plates with Lipid-DNA complexes (500 ng plasmid, 3 μl Lipofectamine LTX, 1 μl Plus Reagent (Life Technologies)). Cells were collected for analyses 72 h post transfection for all but the Gal4-p65 pre-treatment assays. For EZH2 inhibition, cells were treated with 1 μl 10 μM UNC1999 (Cayman Chemical) at a final dilution of 1.0E-3 μM in i ml growth medium for 72 h. Control cells were treated with 1 μl of DMSO (vehicle) per 1 ml media. Cells were grown in fresh UNC1999-free, DMSO-free medium for 24 h prior to Luciferase assays (described below) or ChIP-qPCR (Supporting Information).

### Luciferase Assays

Transfection efficiencies for Gal4-p65-expressing cells were determined by measuring the percent of live, mCherry-positive cells using the BD Accuri C6 Flow Cytometer and accompanying software (BD Biosciences) as previously described^30^. Luciferase signal from 100 μl of cells was normalized by the number of live cells, which was determined by flow cytometry (using side- and forward-scatter gating) of 20 μl from each sample. Luciferase assays were performed in triplicate per experimental or control sample with a Synergy H1 Multi-Mode Reader (Biotek) using the Steady-Luc Firefly HTS Assay Kit (Biotium)^30^. Luciferase activity per cell was calculated as follows:

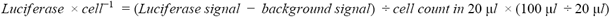

### Deep Sequencing to Determine Cas9 Editing Efficiency

Cells were collected after 72 h of growth following transfection with pU6-(BbsI)_CBh-Cas9-T2A-EGFP (DNASU UnSC00746685) with sgRNA sg025, sg032, sg046, or sg048.^10^ Transfection efficiency was determined using flow cytometry to measure EGFP signal as previously described.^30^ Genomic DNA was extracted from the remaining 200 μl of cells using the QIAamp DNA Mini Kit (Qiagen) and eluted in 100 μl of nuclease-free water. PCR of gDNA was performed using GoTaq 2x Mastermix (Promega) using primers P198/198 (see Table S2 for sequences) [95°C for 10 min; 34x (95 °C, 30 sec; 58 °C, 30 sec; 72 °C, 20 sec.)]. Nested PCR was performed by diluting 2 μl of PCR product into 500 μl of water and Phusion High-Fidelity DNA Polymerase (Thermo Scientific), using the primers described in Table S1 and S2. DNA was purified from PCR reactions using the Genelute PCR Cleanup Kit (Sigma-Aldrich) and submitted to the Center for Computational and Integrative Biology (CCIB) Core Facility (Massachusetts General Hospital) according to their requirements for sequencing.

### Quantitative PCR of ChIP DNA

Chromatin Immunoprecipitation is described in Supporting Information. Relative quantification of ChIP samples was performed using real-time quantitative PCR and SYBR Green I master mix (Roche) as previously described.10 To adjust for input dilution, log2(20) was subtracted from input Cp values. Enrichment of H3K27me3 and H3K4me3 (%IP DNA bound of input) was calculated as 100 × 2^(Cp_input_ - Cp_ip_). Luciferase primer sequences were as follows: P360 5‘-CGGCGCCATTCTATCCTCTA-3’ (forward); P361 5‘-ATTCCGCGTACGTGATGTTC-3’ (reverse). TBP primer sequences: P363 5‘-CAGGGGTTCAGTGAGGTCG-3’ (forward); P364 5‘-CCCTGGGTCACTGCAAAGAT-3’ (reverse).

### Statistical Analyses

The percentage of edited reads was normalized to transfection efficiency for each sample. Averages and standard deviations were calculated for each of 3 biological replicates. To analyze ChIP-qPCR assay data, averages and standard deviations were calculated for each of 3 replicate IPs from a single chromatin preparation. For each cell/ treatment data set, %IP DNA for *luciferase* and *TBP* were normalized to %IP DNA for *TBP*. The differences of means to compare cell/ treatment data sets for both CRISPR editing and ChIP-qPCR experiments were calculated using the two sample, one-tailed Student’s t test: for *p* < 0.05, 95% confidence with 2 degrees of freedom and a test statistic of t_(0.05,2)_= 2.920; for *p* <0.01, 99% confidence with 2 degrees of freedom and a test statistic of t_(0.01,2)_ = 6.965.

## RESULTS

### The fusion activator Gal4-p65 alters the transcriptional state and histone modifications at PRC-repressed chromatin

Our previous work demonstrated that a fusion transcriptional activator composed of the Gal4 DNA binding domain and the p65 protein (from NF-kB) can activate gene expression at a PcG-silenced *luciferase* gene.^10^ P65 is known to interact with several transcriptional activators that potentially counteract PcG-mediated siencing, including histone acetyltransferases (CREBBP and EP300), and H3K4 methyltransferases (EHMT1 and SETD7) (STRING database^31^). We asked whether p65 targeting near the promoter and its presumed recruitment of endogenous factors could induce sustained transcriptional activation and chromatin remodeling at the PcG-silenced *luciferase* transgene.

We used Gal4-EED/luc cells as a model system for PRC-mediated closed chromatin. Upon treatment with doxycycline (dox), the cells express a Gal4-embryonic ectoderm development (EED) fusion protein which binds to a Gal4 enhancer sequence (UAS) upstream of the *thymidine kinase* (*Tk*) core promoter of a *luciferase* transgene. EED recruits PcG proteins (Fig. 1), resulting in accumulation of the H3K27me3 silencing mark, the PRC1 complex, and *luciferase* silencing.^10,29^ As a control for the open chromatin state, we used the Luc14 cell line which does not contain Gal4-EED. The absence of the silencing mark H3K27me3 at *luciferase* in Luc14 cells was confirmed in Daer et al. 2017. Since the *luciferase* transgene had been generated by random chromosomal integration^29^, the distance between endogenous DNA and the PRC nucleation site (UAS) was unknown. Using a series of PCRs with overlapping amplicons, we estimated that the transgene-genome boundary is 2050 to 2457 bp upstream of the Polycomb chromatin nucleation site at *UAS* (Fig. S1). This result, together with previous studies of the dynamics of the system, suggest that *UAS-Tk-luciferase* not significantly influenced by surrounding endogenous chromatin.

We measured the change in *luciferase* expression, the silencing-associated H3K27me3 mark, and the activation-associated H3K4me3 mark in cells expressing the Gal4-p65 activator. We detected strong and sustained re-activation of *luciferase* at 3 days and 9 days following transfection with Gal4-p65-expressing plasmids (Fig. 2A). We also detected a significant reduction in the amount of H3K27me3 (*p* < 0.05) (Fig. 2B). To determine whether p65 targeting results in addition of active chromatin modifications, we measured enrichment of H3K4me3. We observed less H3K4me3 in the silenced state compared to Luc14 (*p* = 0.06) (Fig. 2B) and a significant increase of H3K4me3 in the reopened state versus the silenced state (*p* < 0.01).

**Figure 2.**
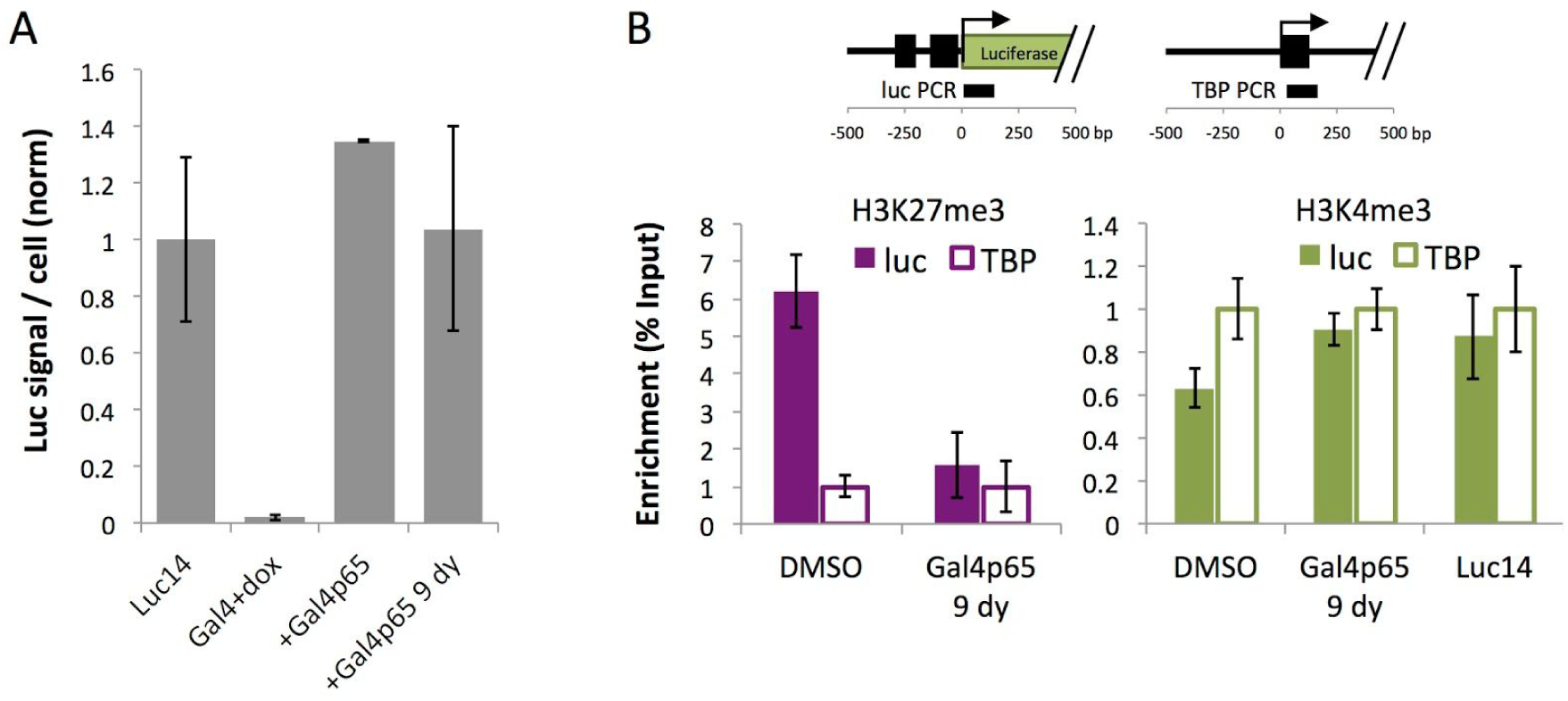
Targeted transcriptional activation by Gal4-p65 leads to increased *luciferase* expression and histone modification. (A) Luciferase activity assays were used to determine *luciferase* expression in Luc14 (open chromatin), dox-treated Gal4-EED/luc cells (Gal4+dox, closed chromatin), and Gal4-p65-expressing Gal4+dox cells either 72 h after transfection (+Gal4p65) or 216 h after transfection (+Gal4p65 9 dy). Each value from 3 biological replicates was normalized to the mean Luc14 value (bars, mean values; error bars, standard deviation). (B) Chromatin immunoprecipitation followed by quantitative PCR (ChIP-qPCR) was used to measure H3K27me3 enrichment at *luciferase* after treatment with UNC1999 compared to the vehicle control. A constitutively active control locus, TATA-binding protein (TBP), was used as a negative or positive control for H3K27me3 or H3K4me3 enrichment, respectively. Each value from 3 replicate pulldowns from a single chromatin prep was normalized to the mean TBP value (bars, mean values; error bars, standard deviation).

Interestingly, our data reveal that PRC-repressed *luciferase* is both H3K27me3-positive and H3K4me3-positive (Fig. 2B). A previous study of the Gal4-EED/luc cell line^29^ showed that PRC accumulation leads to a reduction in, but not a complete disappearance of H3K4me3 at *luciferase*. This result is consistent with the appearance of “bivalent” chromatin at poised, inactive genes^32^ and suggests that silenced *UAS-Tk-luciferase* may be a useful model to study the impact of bivalent chromatin on CRISPR activity. Still, H3K4me3 levels were significantly higher after Gal4-p65-induced *luciferase* activation, suggesting that a greater proportion of H3K4me3 to H3K27me3 is associated active expression.

### Chromatin de-silencing by Gal4-p65 restores CRISPR activity at a distal site

Gal4-p65-induced activation of *luciferase* and the accompanying changes in chromatin structure (Fig. 2B) led us to hypothesize that local chromatin remodeling via Gal4-p65 would enhance Cas9-mediated editing at a site that was previously blocked by closed chromatin. In our 2017 study, we tested this idea by co-transfecting *luciferase*-silenced HEK293 cells with a Gal4-p65-expressing and Cas9/gRNA-expressing plasmids.^10^ *Luciferase* became activated, suggesting a transcriptionally permissive state, but editing efficiency was not significantly higher at the target site we tested, sg034. Also in the previous study, we observed a decrease in editing efficiency at sg034 in the open chromatin state that was hyper-activated by Gal4-p65 suggesting competition between Gal4-p65 and Cas9/gRNA might contribute to reduction in Cas9 activity.

In the current study, we carried out the same treatment (co-expression of Gal4-p65 with CRISPR plasmids) at sites located in the same 1.1 kb region as sg034: sg046, 32, 25, and 48 (Fig. 2B, Table S2). Sg046, 32, and 48 were chosen to represent sites where Cas9 activity is inhibited by closed chromatin.^10^ These sites also represent varying distances from the activator binding site: sg046 and 32 are proximal to the Gal4 UAS sequence (within 300 bp) while sg048 is distal (>1000 bp downstream of the TSS). Sg025 shows no change in editing efficiency within closed chromatin^10^ and was therefore used here as a control for uninhibited CRISPR activity. At 72 hours post-transfection, we collected the treated cells and assayed for Cas9 editing using deep sequencing to quantify the proportions of insertion and deletion (INDEL) variants generated by non-homologous end-joining (NHEJ) repair. We detected editing efficiencies at or below closed chromatin levels for sg032, 25, and 48 or sg046, respectively, (Fig. 2B, Closed +Gal4-p65).^10^ The most frequent INDELs showed no obvious change in position relative the predicted cut sites (Fig. 2C), suggesting no change in Cas9 specificity or NHEJ products.

To reconcile the opposing effects of Gal4-p65 targeting, i.e. induction of an active chromatin state versus inhibition of Cas9‘s access to DNA, we asked whether transient Gal4-p65 expression followed by a recovery period to allow for depletion of Gal4-p65 (referred to here as “pre-treatment”) would generate a Cas9-accessible state. Having established that transcription and activation-associated histone modifications were sustained 9 days after initial overexpression of Gal4-p65 (Fig. 2), we set out to determine CRISPR activity at *luciferase* after Gal4-p65 pre-treatment, compared to co-treatment with Gal4-p65 and CRISPR (Fig. 3A). For pre-treatment, we transfected the Gal4-p65 plasmid alone and then one of the four Cas9/gRNA plasmids 9 days later (2 cell passages). As with co-treatment, we did not observe significant differences in editing efficiency at the control site sg025. At one proximal site (sg032) and the distal site (sg048), the mean editing efficiency approached full recovery (open chromatin levels). The most significant enhancement of editing was observed at the distal site sg048 (*p* < 0.005, Closed +Gal4-p65, 9 day recovery versus closed chromatin). The target site closest to the heterochromatin nucleation site and the TSS, sg046, showed no significant increase in editing compared to the closed chromatin control sample. Overall, these results suggest that initial, transient overexpression results in Gal4-p65 accumulation and activator recruitment that inhibits Cas9 activity by blocking access of Cas9/gRNA to the target sites. This is followed by gradual remodeling of chromatin into a state that enhances Cas9-mediated editing at sites downstream of the TSS.

**Figure 3.**
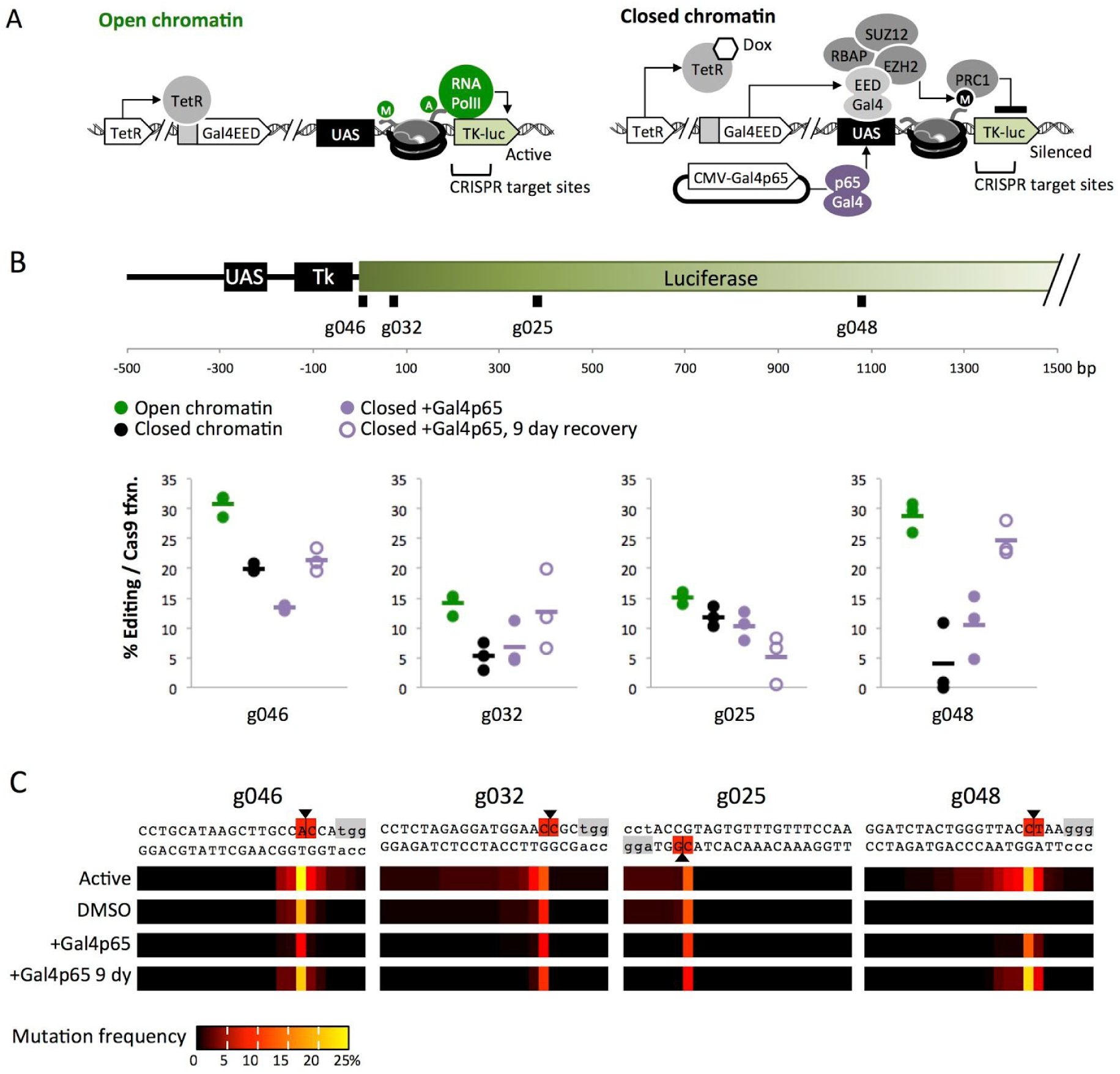
Pre-treatment with a transcriptional activator improves Cas9 editing at closed chromatin. (A) Summary of the experimental approach. Open chromatin is characterized by transcription of *luciferase* and absence of the silencing mark histone 3 lysine 27 trimethylation (H3K27me3).^10^ Closed chromatin is induced upon addition of doxycycline (dox) and is characterized by reduced expression at *luciferase* (Fig. 2A) and accumulation of H3K27me3 (Fig. 2B and Daer et al. 2017). Dox-treated Gal4-EED/luc cells were transfected with the CMV-Gal4-p65 plasmid, which results in long-term activation of *luciferase* expression and local chromatin remodeling (Fig. 2B). (B) Comparison of Cas9 editing efficiencies at each sgRNA target site for different chromatin states. The luciferase map, open chromatin, and closed chromatin samples are described in Figure 3. “Closed +Gal4-p65” (purple filled) are Gal4-EED/luc+dox silenced cells that were co-transfected with CMV-Gal4-p65 plasmid and Cas9/sgRNA expressing plasmids with sgRNAs targeting sg046, 32, 25, and 48 at 24 hours after dox removal. “Closed +Gal4-p65, 9 day recovery” (purple outlined) are Gal4+dox silenced cells transfected with only CMV-Gal4-p65 plasmid at 24 hours post dox removal, then cultured for 216 hours (9 days) and then transfected with Cas9/sgRNA plasmids. For all samples, editing efficiencies were measured after 72 hours of Cas9/sgRNA expression. Dots, individual replicates; bars, means of three biological replicates. (C) Heat maps indicate the frequency at which each DNA base position was affected by an insertion or deletion (INDEL). One representative biological replicate is shown for each condition.

### Depletion of H3K27me3 via EZH2 inhibition is not sufficient to generate an active expression state

Next, we investigated the contribution of H3K27me3 demethylation to de-silencing of PRC-repressed *luciferase* and restoration of CRISPR-mediated editing. UNC1999 is a small molecule drug that binds to the active site of the methyltransferase Enhancer of zeste 2 (EZH2) to halt the methylation at H3K27me3, a modification that supports PRC formation and a closed chromatin state.^33^ UNC1999 is a potent inhibitor that has been used for complete depletion of H3K27me3 in MCF7 and HEK293 cells.^33^ We induced repressive chromatin at *luciferase* with dox-supplemented media for 96 hours. At 120 hours, we cultured the cells in a sub-lethal dose of UNC1999 (1.0E-3 μM) for 120 hours. At 264 hours, we measured Luciferase activity and performed chromatin immunoprecipitation followed by quantitative PCR (ChIP-qPCR) to determine H3K27me3 enrichment.

While UNC1999 did not significantly increase Luciferase activity (Fig. 4A), we saw a significant decrease in H3K27me3 compared to the DMSO control (*p* < 0.01) (Fig. 4B). We conclude that at 1.0E-3 μM, UNC1999 depletes H3K27me3 from the silenced transgene but does not stimulate transcriptional activation under the conditions tested here. This finding is consistent with previous work where removal of the H3K27me3 silencing mark alone did not activate gene expression in embryonic stem cells,^34^ and DNA methylation and gene silencing persisted after depletion of EZH2 in cancer cells.^35^ The effect of UNC1999 allowed us to investigate the impact of H3K27 modification on CRISPR activity independently of the elevated expression that was observed with transcriptional activation via Gal4-p65.

**Figure 4.**
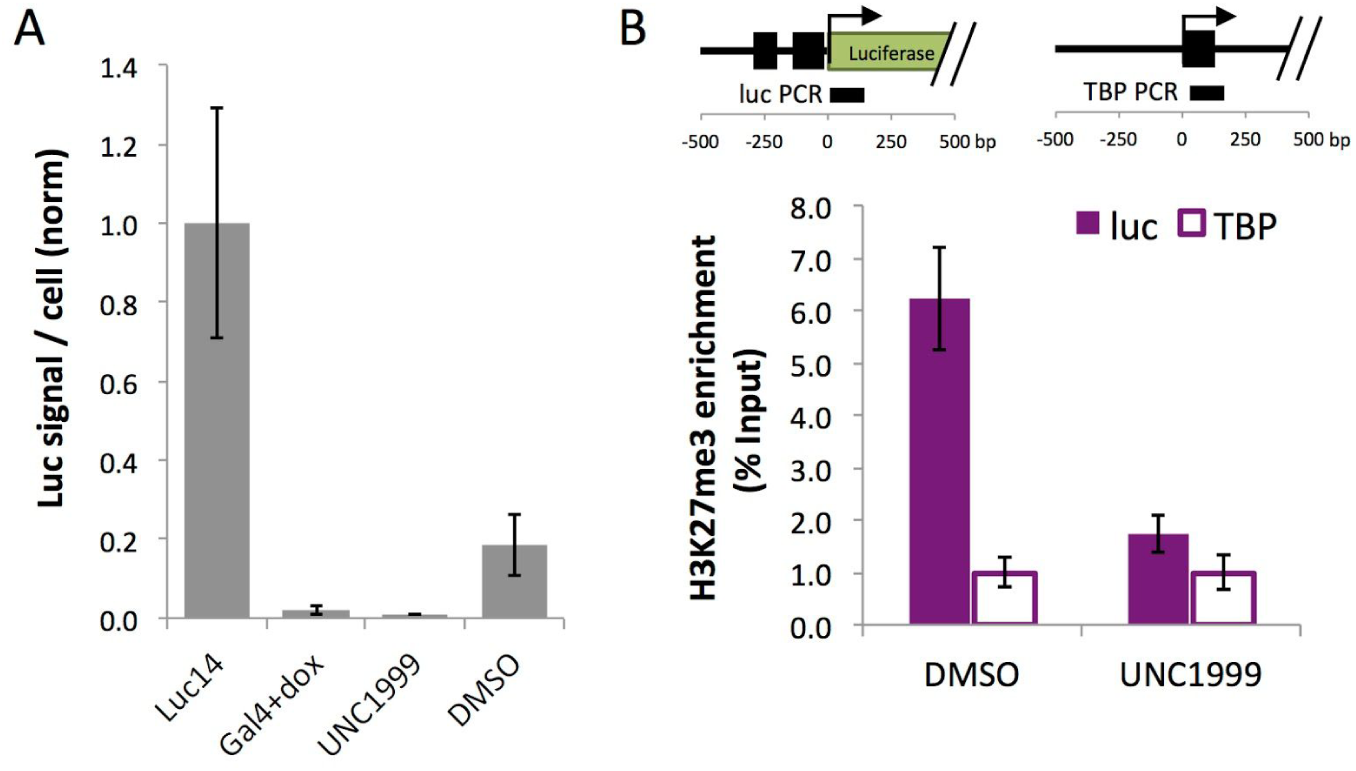
UNC1999 depletes histone 3 lysine 27 trimethylation (H3K27me3). (A) Luciferase activity assays were used to determine *luciferase* expression in Luc14 (open chromatin), dox-treated Gal4-EED/luc cells (Gal4+dox, closed chromatin), and Gal4+dox cells treated with an EZH2 inhibitor (UNC1999) or the vehicle control (DMSO). Each value from 3 biological replicates was normalized to the mean Luc14 value (bars, mean values; error bars, standard deviation). (B) ChIP-qPCR was performed and data were analyzed as described in Figure 2. DMSO sample data is the same as in Figure 2.

### Depletion of H3K27me3 by inhibition of EZH2 is not sufficient to restore CRISPR activity in PRC-repressed chromatin

Next, we investigated whether removal of H3K27me3 via inhibition of EZH2 (Fig. 5A) was sufficient to improve Cas9 editing. We measured Cas9 activity in cells that were subjected to the same DMSO and UNC1999 treatments we used for the expression and ChIP-qPCR assays. At 24 hours after DMSO or UNC1999 removal, we transfected cells with plasmids expressing Cas9/sgRNA that targetes site sg046, 32, 25, or 48 at *luciferase* (Fig. 5B). At 72 hours post-transfection, we collected the treated cells and assayed for Cas9 editing using deep sequencing. UNC1999 treatment did not restore Cas9 editing at any of the four target sites tested. Furthermore, at two target sites, treatment with UNC1999 showed an even greater inhibition than in the vehicle control (sg046 *p* < 0.001 and sg025 *p* < 0.01). Since inhibition of endogenous EZH2 has a global effect on genomic chromatin, it is difficult to interpret this result. The position of the most frequently edited sites (the nucleotide immediately adjacent the predicted cut site) was the same under all conditions (Fig. 5C), suggesting that treatment with UNC1999 did not alter the pattern of editing. We conclude that UNC1999 treatment under these conditions does not improve Cas9-mediated editing at sites in PRC-repressed chromatin.

**Figure 5.**
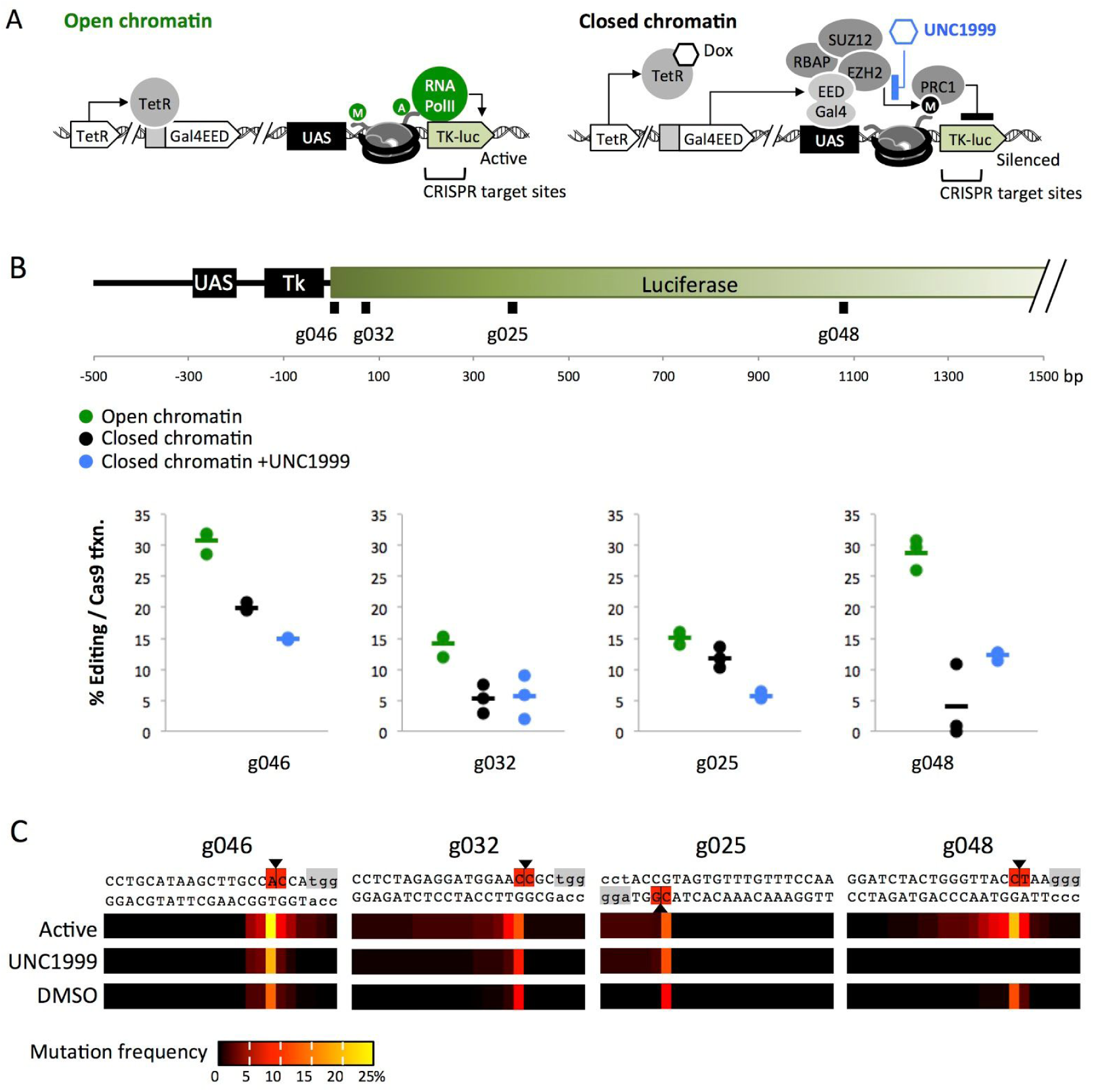
Treatment with UNC1999 does not restore Cas9-mediated editing. (A) Overview of the experimental approach. Chromatin states are shown as described in Figure 3. Treatment with UNC1999 (blue hexagon) inhibits Enhancer of zeste 2 (EZH2), resulting in the loss of H3K27me3 (Fig. 4B and Konze et al. 2013). (B) Map of the *luciferase* transgene and the sgRNA target sites. Charts show Cas9 editing efficiencies (deep sequencing data) at each target site for three different conditions. “Open chromatin” (green) represents untreated Luc14 cells transfected with Cas9/sgRNA expressing plasmids with sgRNAs targeting sg046, 32, 25, and 48. “Closed chromatin” (black) and “Closed chromatin +UNC1999” (blue) are Gal4-EED/luc+dox silenced cells treated with either the vehicle (DMSO) or UNC1999, respectively. For all samples, editing efficiencies were measured 72 hours post transfection with Cas9/sgRNA plasmids. Dots, individual replicates; bars, means of three biological replicates. (C) Heat maps of the mutation frequency for a representative biological replicate. “Open chromatin” and “Closed chromatin” values are the same data presented in Figure 3.

## DISCUSSION

We determined the impact of de-silenced and H3K27me3-depleted Polycomb-enriched chromatin within a 1000 bp chromosomal region in a human cell line. We targeted spCas9 to four sites along a *luciferase* transgene within ectopic PRC-associated chromatin. Three sites showed inhibition by heterochromatin (sg046, 32, and 48) while one site (sg025) was unaffected. We confirmed our observation from previous work that strong transcriptional activation driven by Gal4-p65 initially inhibits Cas9 activity.^10^ A recovery period of 9 days following Gal4-p65 expression (pre-treatment) lead to sustained activation of luciferase, loss of H3K27me3, an increase in H3K4me3, and enhancement of Cas9/gRNA activity at two sites. The most significant enhancement was observed for a site ∼1000 bp downstream of the activator target site.

While UNC1999 effectively removed H3K27me3 from the *luciferase* transgene (Fig. 4B), it was not sufficient to induce gene activation (Fig. 4A) or enhance Cas9 editing at the sites tested (Fig. 5B and C). This is consistent with previous observations where removal of H3K27me3 did not result in gene activation^34^ or remodeling of chromatin to an open state; gene silencing and other repressive modifications can persist after the loss of PcG proteins.^35^ siRNA-mediated knockdown of the non-enzymatic PRC protein Suz12 showed modest enhancement of CRISPR activity in our previous study^30^, as well as a small increase in *luciferase* expression. Along with the results from our Gal4-p65 experiments, these observations suggest a role for gene expression or expression-associated chromatin modifications in generating the Cas9/gRNA-accessible state. Expression-associated chromatin modifications may work in concert with or independently of H3K27me3 depletion to open repressed chromatin. Specific inhibitors of epigenetic enzymes other than the compound investigated here warrant further investigation. These could be used to interrogate the role of other chromatin features, including DNA methylation and histone deacetylation, in modulating access of DNA by CRISPR.

We demonstrated that targeted remodeling using a strong activator (p65) enhances Cas9-mediated editing roughly 1000 bp downstream from the Gal4 binding site (UAS) of a model transgene, *luciferase*. Targeted hyper-activation can generate a Cas9-accessible state in the ectopic chromatin model system used here, under specific conditions. We confirmed our previous observation that high-transcriptional activity can impede Cas9 editing possibly through steric hindrance and competition with the high occupancy of the UAS site with activator proteins and RNA PolII. Co-treatment with the Gal4-p65 activator showed no improvement in editing and, at one site, significant inhibition compared to the silenced state (Fig. 3B and C). In contrast, pre-treatment with the activator enhanced Cas9 editing near open-chromatin levels at target sites previously inhibited by closed chromatin (Fig. 3B and C). Pre-treatment also led to depletion of the H3K27me3 silencing mark, accumulation of H3K4me3 activating marks, and gene activation (Fig. 2). This suggests that chromatin remodelers recruited by Gal4-p65, i.e. histone acetyltransferases (CREBBP and EP300), and H3K4 methyltransferases (EHMT1 and SETD7), or transcriptional elongation remove or displace nucleosomes or the silencing marks.

To make these findings more broadly applicable for CRISPR enhancement, the Gal4 binding domain could be replaced with other customizable DNA binding modules such as dCas9/sgRNA, zinc fingers (ZF) or transcription activator-like (TAL) effectors and be targeted to accessible DNA sites up- or down-stream of the desired endogenous target gene.^28^ However, if enhanced CRISPR editing relies on transcriptional elongation, the effect of p65 would require proximity to a transcription initiation complex. Furthermore, transcriptional activation and expression of certain gene products during the DNA engineering process could destabilize cell phenotypes and make the development of engineered cells extremely difficult. Therefore, effective fusion-chromatin remodelers that function independently of promoter-proximal-p65 to generate a CRISPR-accessible state could provide useful options. So far, Barkal et al. demonstrated that non-enzymatic dCas9 increased access of DNA within 100 bp of the dCas9 binding site, while having a neutral effect on gene expression^17,18^. Our study is a significant step towards manipulating chromatin across broader regions to enhance the efficacy of CRISPR-mediated genome editing.

## ACKNOWLEDGEMENTS

This work was supported by NSF CBET (1404084). RMD was supported by the ARCS Foundation. KAH was supported by NIH NCI (K01 CA188164). The authors thank the Center for Computational and Integrative Biology (CCIB) Core Facility (Massachusetts General Hospital) for providing CRISPR sequencing services. The authors thank J. Steel (DNASU) for the use of their Tapestation. TBP primers were a generous gift from J. Cutts (Brafman Lab at ASU)and the Luc14 and GAL4-EED cell lines were a generous gift from K. Hansen. The authors thank J. Daer for critical review of this manuscript.

## AUTHOR CONTRIBUTIONS

R.D. developed the assays, performed DNA construction, transfections, preparation of samples for deep sequencing, and ChIP-qPCR assays. C.M.B. performed luciferase assays and assisted with ChIP-qPCR. K.A.H. oversaw the experiments, assisted with data analysis, and created the figures. All co-authors All co-authors have reviewed and approved of the manuscript prior to submission.

